# dgiLIT: A Method for Prioritization and AI Curation of Drug-Gene Interactions

**DOI:** 10.64898/2026.01.16.699733

**Authors:** Matthew Cannon, Anastasia Bratulin, James Stevenson, Kathryn Perry, Adam Coffman, Susanna Kiwala, Lars Schimmelpfennig, Heather Costello, Joshua F McMichael, Malachi Griffith, Obi L. Griffith, Alex H. Wagner

## Abstract

**IMPORTANCE:** The Drug-Gene Interaction Database (DGIdb) has a long history of driving hypothesis generation for biomedical research through the careful curation of drug-gene interaction data from primary and secondary sources with supporting literature. Recent advances in large-language model (LLM) and artificial intelligence (AI) technologies have enabled new paradigms for knowledge extraction and biocuration. The accelerating growth of biomedical literature presents a significant challenge for maintaining up-to-date interaction data. With more than 38 million citations indexed in PubMed alone, new strategies must evolve to identify and incorporate new interaction data into DGIdb.

**OBJECTIVE:** Identify new cost-effective AI curation strategies for incorporating new drug-gene interactions into DGIdb.

**METHODS:** We present a methodology that leverages deterministic natural language processing techniques, existing harmonization frameworks, and AI-assisted curation to systematically narrow the literature space and identify new drug-gene interactions from published studies for inclusion in DGIdb.

**RESULTS:** We demonstrate the use of lemmatization to prioritize a set of 100 abstracts containing high amounts of interaction words for downstream AI curation. From our set of abstracts, we were then able to identify 137 drug-gene interactions via an AI curation task, with 121 (88.3%) of these interactions being completely novel to DGIdb. A human expert evaluator reviewed this interaction set and was able to validate 134 of 137 (97.8%) interactions as being valid based on the text provided.

**CONCLUSION:** Taken together, our results highlight a promising, cost-effective method of ingesting new interactions into DGIdb.

## INTRODUCTION

The Drug-Gene Interaction database (DGIdb) is a resource that acts as a centralized aggregator for drugs, genes, interactions, and category annotations from 45 different sources. The central mission of our resource has been to harmonize drug-gene interaction data from disparate sources in an effort to elucidate unique insights and new hypotheses for the druggable genome that might not otherwise be found. DGIdb has a long history of driving hypothesis generation for biomedical and biochemical research and is widely used, garnering 3,263 citations from the research community spanning its 12 year publication history^1–5^. By importing and performing rigorous grouping methods on drug and gene concepts, DGIdb provides a toolkit for biomedical researchers to evaluate druggability of genes and therapeutics across different contexts.

The drug-gene interaction data available from our resource and others would not be possible without significant curation efforts. The majority of biomedical knowledge originates from peer-reviewed, published literature and often exists as unstructured text that requires sustained human curatorial labor^6,7^. Notably, one illustrative example of data richness can be found from the original *Hopkins and Groom (2002)* and *Russ and Lampel (2005)* publications describing not only the concept of a ‘druggable genome’, but also containing over 399 molecular targets and 2917 druggable genes, respectively^8,9^. With these quantities of data originating from only two publications, there is likely a substantial amount of potential therapeutic interaction data hidden amongst the more than 38 million published articles indexed by PubMed^10^.

The rich yet computationally inaccessible knowledge in publications has been a driving motivation for the development of many text mining, text annotation, and knowledge extraction efforts^11,12^. Foundational works, such as those in *Ren et al (2018)* and *Singhal et al (2016)*, presented powerful methods for extraction of published knowledge for biomedical curation^13,14^. In recent years, the advent of large language models (LLM) and artificial intelligence (AI) technologies have enabled further advances in knowledge extraction from biomedical corpora^15^. Notably, these advances include the introduction of LLMs pre-trained specifically for biomedical tasks, such as BioBERT and SciBERT (among others)^16–19^. While these models demonstrated state-of-the-art performance with high accuracy and recall on specific benchmarked tasks (e.g. entity recognition, relation extraction, and text summarization), their applications for real-world clinical workflows still remain tenuous due largely to their propensity for producing hallucinations^20–22^. The stakes are high in the medical field, and any potential for misinformation represents an unacceptable risk to a patient’s wellbeing. Thus, applications of LLMs in the medical field seem to have the most success and potential for benefits when deployed in low-risk, assistive knowledge extraction roles^23–26^. In these scenarios, the previously time-intensive knowledge discovery phase is reduced, and human experts can instead spend their available time synthesizing knowledge for their respective clinical roles.

The gap between published biomedical research and its availability as computable, clinically applicable medical knowledge presents a unique opportunity for our resource to apply emerging LLM technologies. With AI-centric approaches proving effective at knowledge extraction tasks, there is ample room to apply LLMs towards extraction of new drug-gene interactions from literature. However, the volume of available literature renders brute force AI methodologies costly both in computational time and financial value. Similarly, the previously discussed hallucination-prone nature of LLMs supports the need for human guard rails to ensure the validity of AI extracted drug-gene interaction data.

To address this need, we present the drug-gene interaction literature integration tool (dgiLIT) as an LLM-assistive toolkit to curate new drug-gene interactions into DGIdb. dgiLIT utilizes a combined approach of basic natural language processing techniques, existing harmonization technologies, and AI-assisted curation to retrieve, prioritize, and curate drug-gene interactions from published literature. In this report, we describe how our tool prioritizes gene-specific literature by the number of contained interaction words and pre-processes them for a downstream curation task by an AI agent. Once pre-processed, interactions are extracted via LLM from text and validated by a combination of data harmonization and human review. We demonstrate that the use of dgiLIT can produce high quality, human validated interactions for inclusion in DGIdb as a new, searchable layer of interaction data for downstream applications.

## MAIN TEXT

### Curation of Literature using dgiLIT

In an effort to curate drug-gene interaction data from the “data ocean” of biomedical literature into DGIdb, we developed the Drug-Gene Interaction Literature Integration Tool (dgiLIT). This toolkit provides a four-step process to enable prioritization and subsequent AI-assisted curation of drug-gene interactions from published biomedical literature in PubMed **(Figure 1)**. Identifying the most relevant subset of publications for a given query remains a significant challenge, even with modern search capabilities^10^. To address this, dgiLIT uses the PubTator3 annotation set^27,28^ to subset literature for PMIDs that contain a user’s key genes of interest. While our chosen selection method is imperfect, it allows us to use the state-of-the-art set of generated annotations from PubTator3 to restrict our search to only those related to a particular gene of interest. The subset of abstracts is then lemmatized (reduced to their stem word, e.g., interacting becomes interact) and compared against pre-constructed groups of interaction lemmas, or words that would suggest the possibility of a drug-gene interaction. Exact lemma matches from abstracts with these groups are summed in a bag-of-words approach and sorted based on descending aggregate count. Literature with high concordance of relevant interaction lemmas are then processed and scanned via an AI Curation agent for possible drug-gene interactions. AI output is standardized to JSON format. Drugs and genes identified via AI are ontologically grounded with the VICC suite of concept normalizers^29,30^ to programmatically trim erroneous or incorrect entities.

**Fig 1.**
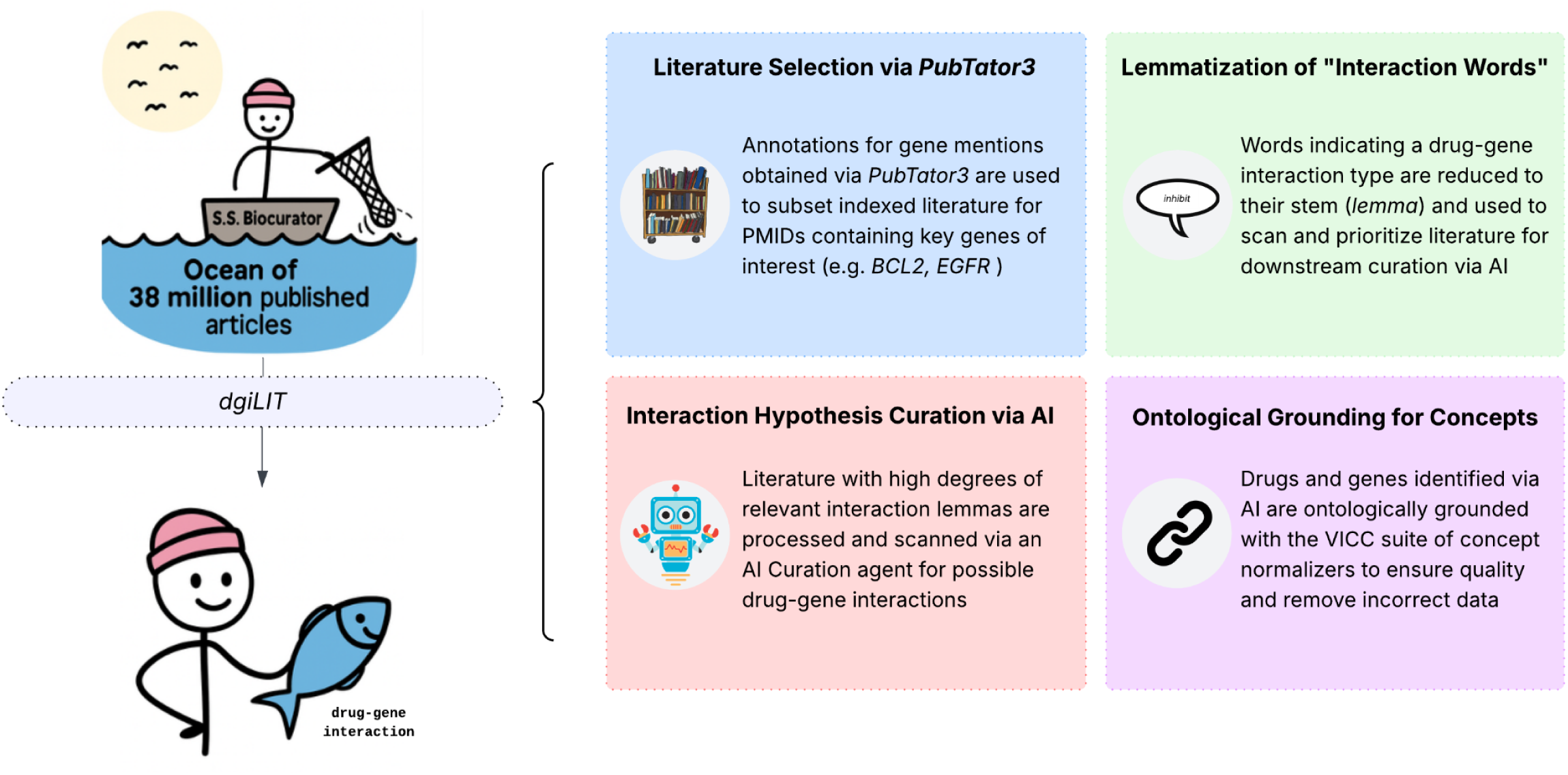
Curating Literature with dgiLIT. The drug-gene interaction literature integration tool (dgiLIT) was developed as a package to enable AI-assisted curation of literature supporting new and existing drug-gene interaction data into DGIdb. Annotations for genes of interest are obtained via *PubTator3* and used to subset literature from indexed PubMed articles. The number of “interaction words” present in abstracts are assessed via lemmatization and are used to further section PMIDs for AI curation. Drug-gene interactions are then curated from selected articles via an AI agent with identified terms normalized via the VICC concept normalization suite.

### Construction of Interaction Lemma Groups

Lemma groups were utilized to capture and quantify “interaction words” across published biomedical abstracts **(Figure 2)**. Lemma group categories were selected from a subset of DGidb interaction type definitions with consideration for common modalities^31^. The categories constructed were: *Gene Regulation, Ligand Binding, Modulation, Pharmacogenomic Signals,* and *Sensitivity and Resistance*. Within each category, words were selected for assignment according to the alignment of common use of the word in biomedical literature and their formal definition^32–39^. For example, the *Ligand Binding* group category contains the lemmas: *ligand, association, recruit, interact, complex, dock, target, bind, affinity, occupy, associate,* and *sequester*. Ontological grounding and definitions for all groups and associated interaction lemmas are provided in **Supplementary Table 1**.

**Fig 2.**
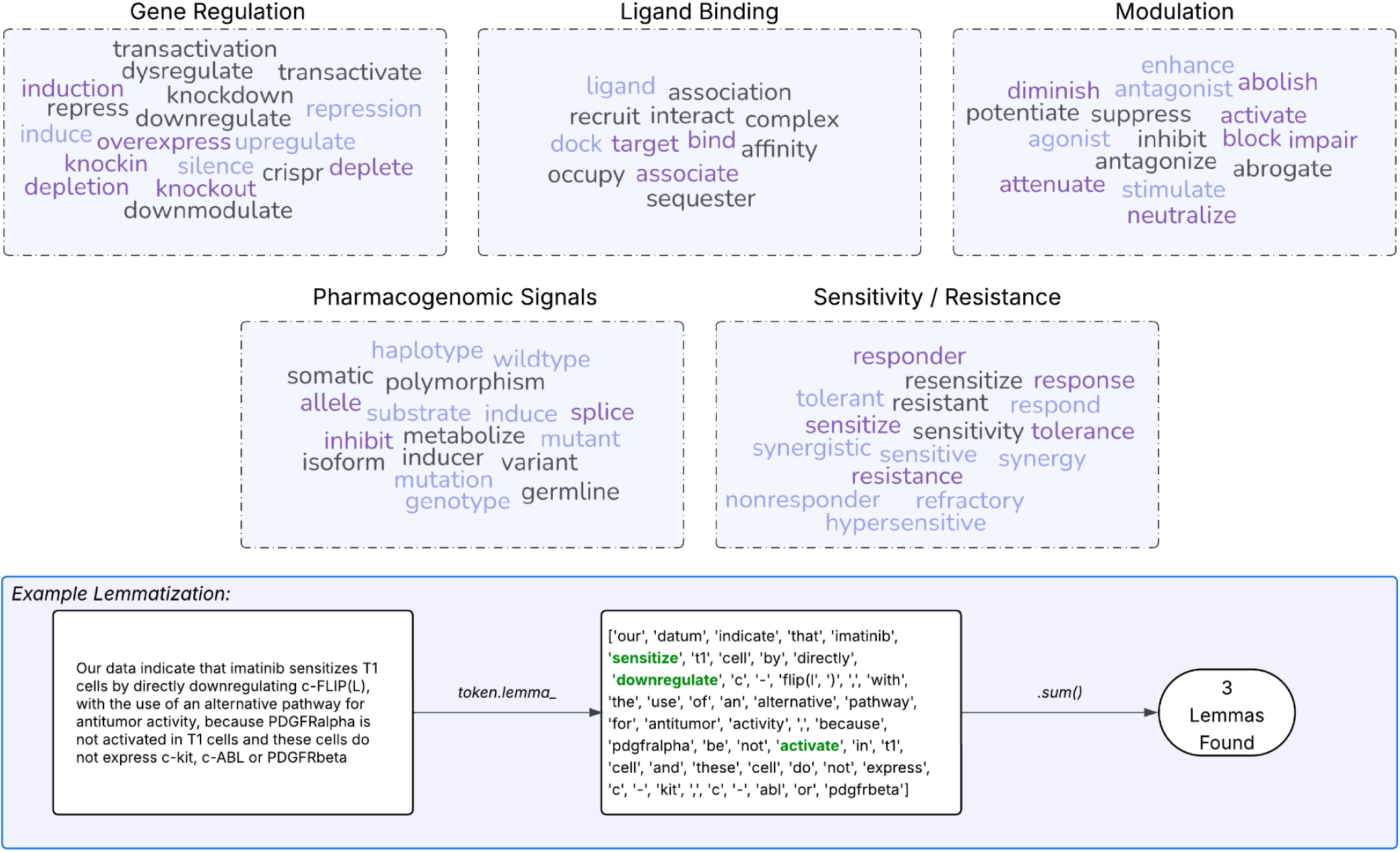
Interaction Lemma Groups. Lemma groups were constructed to capture “interaction words” in literature. Abstract text is lemmatized (e.g. “interacting” becomes “interact”) and exact matches are counted in a bag-of-words approach. Lemma groups were selected from a subset of DGIdb interaction type definitions.

### Bag-of-words Quantification of Interaction Lemmas for Prioritization

Using our described bag-of-words approach, we were able to identify 62,196 interaction-associated lemmas from a set of 14,075 PubTator3-annotated biomedical research articles. These lemmas were subsequently summed and used to naïvely sort data by lemma count. From the total results, the set of 100 published PMIDs containing the highest number of interaction lemmas were selected for downstream AI curation tasks (**Supplementary Table 2**).

### Prompt Creation for AI Curation Tasks

Prompts were programmatically assembled for the set of 100 published PMIDs containing the highest number of interaction lemmas. For each individual publication, a human-designed prompt was created containing a role description, a task description, definitions for *interaction* and *interaction directionality*, a list of drugs to consider for the task, a description of intended JSON output for the task, and a context field (**Figure 3**). With the exception of the list of drugs to consider for the task and the context field, all of these provided fields were identical for every prompt. The list of drugs to consider for the task was obtained from named entity recognition tagging of chemical entities of abstract text using a publically available BioBERT implementation (accessible from the Huggingface ecosystem)^40^. The context field for each prompt contained the abstract text and PMID taken directly from the associated publication. The preprocessed dataframe is available in full from **Supplementary Table 2**.

**Fig 3.**
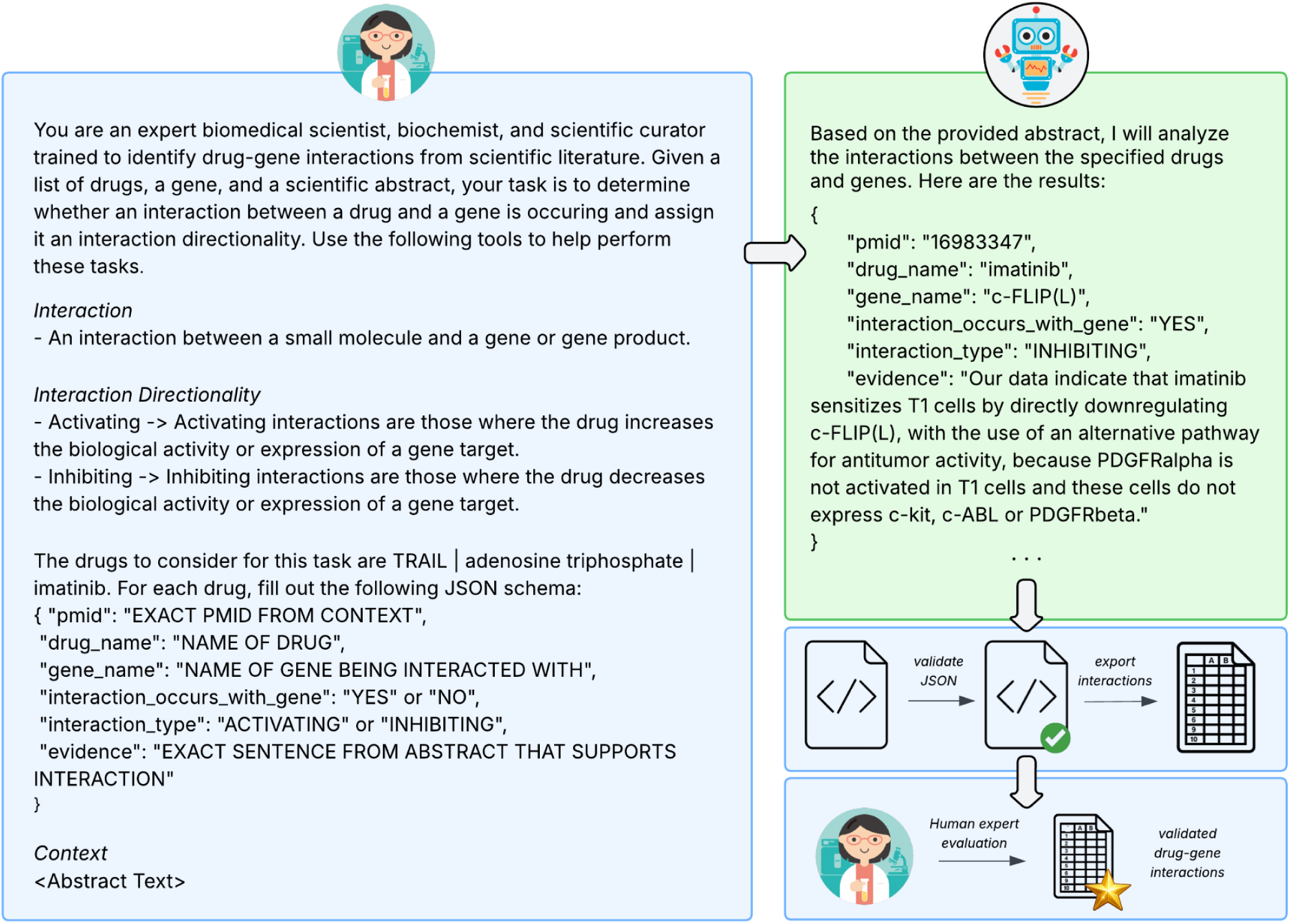
Prompting AI agents for Interaction Curation for DGIdb. Example prompt and response for AI curation of drug-gene interactions for DGIdb. Prompts are created for each abstract in bulk using a basic prompt template containing: a role description, a task description, a toolset description, and a sample output template. The basic prompt is filled in with NER-identified therapeutics and published text as context on an abstract-specific basis. Responses from the AI curation agent are recorded, parsed to extract JSON objects matching the supplied template, and subsequently validated. AI curated interactions are extracted and saved into a *csv* format for human review.

### Interaction Identification via AI assisted curation

A total of 137 interactions were identified via AI curation from our initial set of 100 prioritized publications. Interactions from the response that did not pass initial sanity checks or normalization (e.g. not an interaction, not a valid gene/drug; 40 sanity check failures, 27 normalization failures) were dropped and thus not included in this number. Of these interactions, 121 (88.3%) were completely novel to knowledge already curated within DGIdb (**Figure 4a**). Interestingly, many individual drugs were found multiple times across multiple different curated interactions in our test set, such as the targeted therapeutic venetoclax appearing in 8 interactions (**Figure 4b**). The same pattern was found with genes from this dataset, such as the genes AR and EGFR appearing in 5 interactions each (**Figure 4c**).

**Fig 4.**
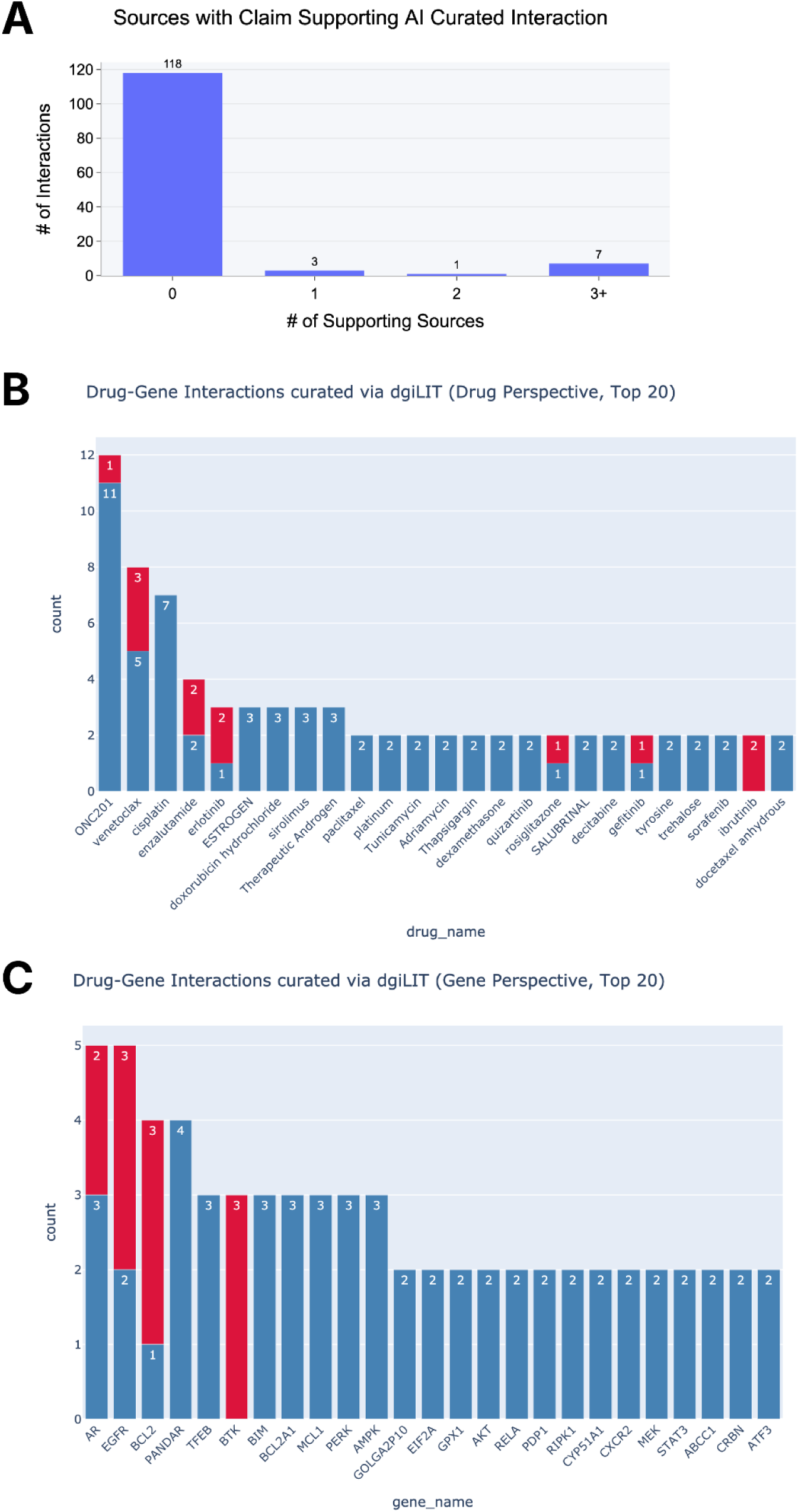
Curation of BCL2 Literature Ingests High-Quality PMIDs and New Interactions into DGIdb. The dgiLIT toolkit was used to select and curate new interactions from 100 PMIDs related to the gene BCL2. (**A**) The number of sources with an interaction claim for unique AI curated interaction. Completely novel interactions have zero claims from other sources. **(B-C)** Most frequently occurring drug and gene partners from the 137 interactions curated via dgiLIT (from drug and gene perspective). Already curated interactions are shown in red.

In addition to the novel interactions identified via AI curation, a number of the individual drug and gene concepts themselves were identified as completely novel knowledge to our resource. Of the 83 unique drug concepts from this dataset, 9 of them were novel to DGIdb (**Table 1**). Similarly, of the 87 unique gene concepts from this dataset, 8 of them were found to be novel (**Table 2**).

**Table 1.**
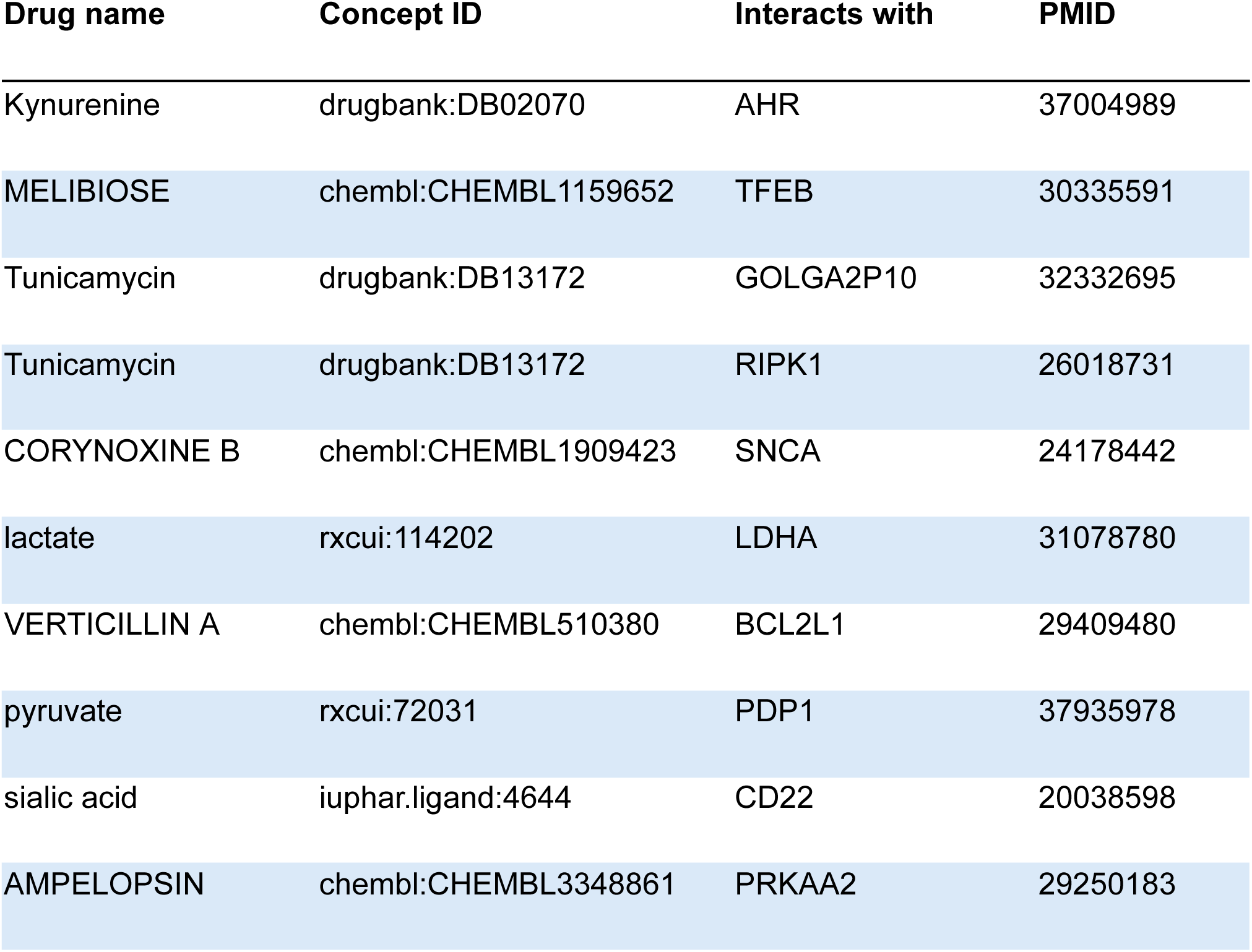
Drug concepts identified via AI Curation agent novel to DGIdb. Drug concepts may appear twice in this list to highlight their involvement in multiple AI curated interactions.

**Table 2.** Gene concepts identified via AI Curation agent novel to DGIdb. Gene concepts may appear twice in this list to highlight their involvement in multiple AI curated interactions.

| Gene name | Concept ID | Interacts with | PMID |
| --- | --- | --- | --- |
| GOLGA2P10 | hgnc:26229 | Tunicamycin | 32332695 |
| GOLGA2P10 | hgnc:26229 | Thapsigargin | 32332695 |
| PANDAR | hgnc:44048 | cisplatin | 30375398 |
| PANDAR | hgnc:44048 | doxorubicin hydrochloride | 30375398 |
| PANDAR | hgnc:44048 | paclitaxel | 30375398 |
| PANDAR | hgnc:44048 | platinum | 30375398 |
| CCDC106 | hgnc:30181 | alanine | 30885251 |
| LINC00173 | hgnc:33791 | cisplatin | 36627670 |
| RASAL2 | hgnc:9874 | platinum | 39890967 |
| EIF2A | hgnc:3254 | ESTROGEN | 30655322 |
| EIF2A | hgnc:3254 | SALUBRINAL | 30655322 |
| TCP1 | hgnc:11655 | Adriamycin | 34750375 |
| MIR375 | hgnc:31868 | enzalutamide | 36042375 |

Classification, citation, and impact profiling of the published articles was performed to assess the credibility of the literature that was used to produce the interaction dataset. Confidence in the relevance of the selected literature towards our resource and its potential for interactions was bolstered by the frequent occurrence of MeSH related to biomedical and drug-related terminology, such as “Apoptosis/drug effects”, “Drug Resistance”, and “Signal transduction/drug effects” (**Supp. Figure 1a**). Similarly, the distributions of journal H-indices and SCImago Journal Rank (SJR) scores demonstrated a high average citation rate and estimated impact factor (**Supp. Figure 1b**).

### Human Expert Evaluation of AI Curated Interaction Evidence

An expert human biocurator with formal training in biocuration and domain expertise in genomic entities and nomenclature was asked to review and validate the output of our AI interaction curation task (**Figure 5**). The human evaluator received the output for each PMID and evaluated the results in a two step process; performing sanity checks on output fields and then performing their own assessment of the validity of the interaction (**Figure 5a**). The raw output of AI interaction curation is provided in **Supplemental Table 3** and the reviewer evaluation form is provided in **Supplemental Table 4**. As part of the sanity check, the evaluator first evaluated whether the drug, gene, and evidence statements provided in the corresponding JSON fields were indeed present in the abstract and not hallucinated. Following this check, the evaluator next validated the interaction based on the results presented in the text and provided their own assessment of the interaction type and directionality (as defined from https://dgidb.org/about/overview/types-and-directionality). When performing this assessment, the evaluator also recorded whether the results presented in the abstract text were sufficient to infer an interaction and, if not sufficient, whether they were able to validate the interaction using additional information from the full-text.

**Figure 5.**
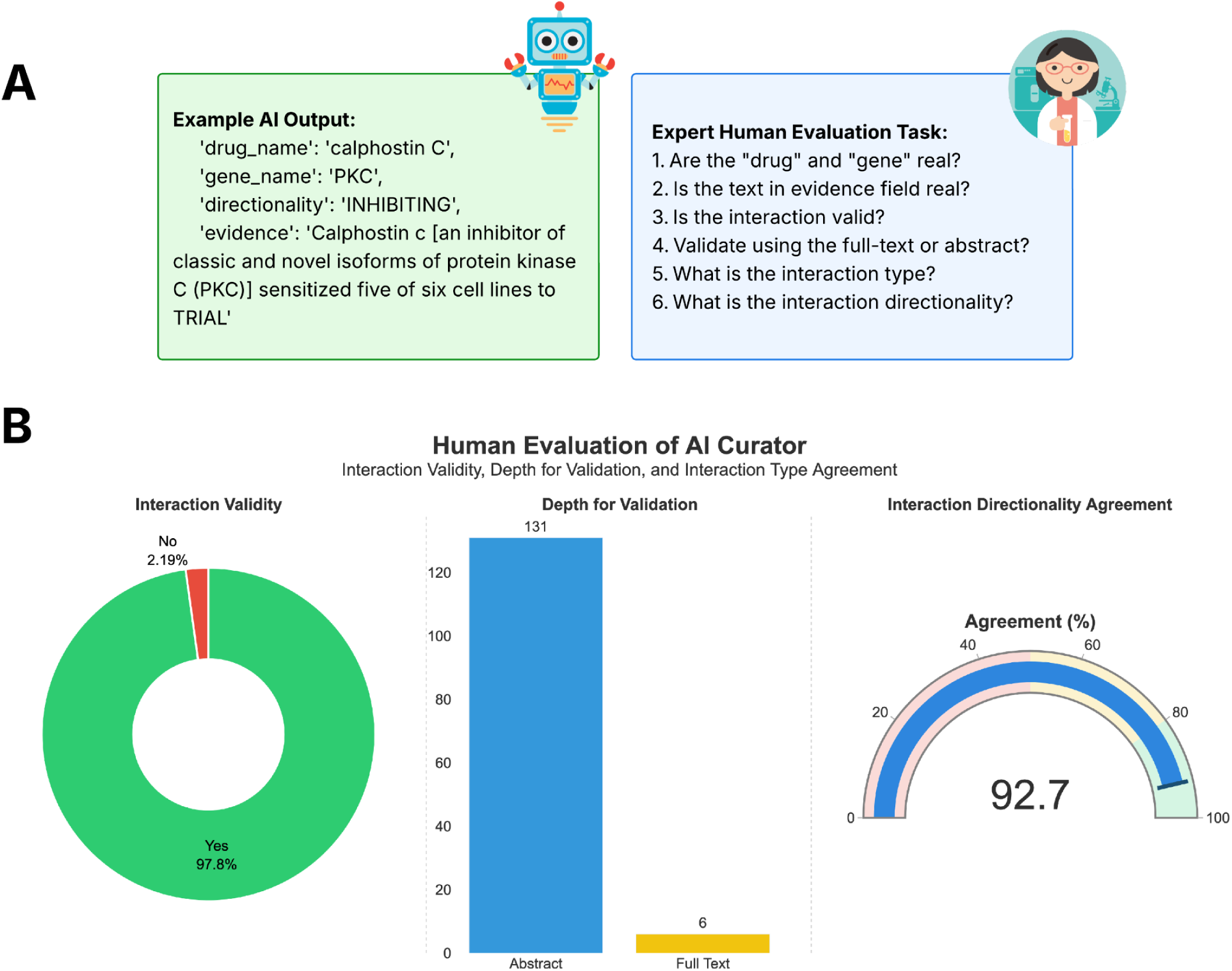
Validation of AI Curated Interactions via Expert Human Review. Overview of workflow and results from human validation tasks. **(A).** Example output of AI curation agent and corresponding validation task for human reviewer. **(B)** Expert review of curated AI output demonstrated a high success rate and agreement for identifying valid interactions and directionality from abstract text alone.

In our assessment of AI curated interactions from BCL2 literature, our human expert was able to validate and assign an interaction type for 134 of 137 (97.8%) AI curated interactions (**Figure 5b, Panel 1**). In this process, the full-text was only required for validation for 6 interactions with the results presented in the abstract sufficient for validation of the remaining 131 interactions (**Figure 5b, Panel 2**). Lastly, our human evaluator showed agreement with the AI assessment of interaction directionality (inhibiting vs activating) for 127 of 137 interactions (92.7%) (**Figure 5b, Panel 3**). The breakdown of interaction types assigned and relative confidence assessments are provided in **Supplementary Figure 2**.

## Discussion

In our report we demonstrated the use of lemmas as a means for prioritizing literature containing interactions. Previous works have already demonstrated the effectiveness of querying for terms directly into a corpus as a means of text mining. In 2019, the method KinderMiner demonstrated this by identifying drug repurposing opportunities from the Europe PMC corpus using a series of transcription factor based queries paired with the key phrase “embryonic stem cell”^41^. Despite its effectiveness in executing this task, the authors take care to note that KinderMiner would ‘almost certainly benefit’ from text normalization and named-entity recognition.

Our methodology, instead, acts as an intentionally simple, cheap heuristic for ‘curation opportunity’ where the more frequently occurring an interaction word, the more likely an interaction. By including normalization of words via lemmatization and scanning for interaction words in this manner, we are prioritizing rapid identification of common representations of interactions in literature^31^. While this method is not foolproof, our results highlight its potential for generating novel interaction hypotheses from previously untapped literature.

We demonstrate LLMs in a role more aligned with modern biocuration norms and more acceptable risk tolerance. Previous work has shown the effectiveness of LLMs at extracting information from medical reports and returning it in a structured format^42,43^. Other work has shown the effectiveness of LLMs at performing relation extraction in a biocuration context^44,45^. In a similar way, we employed our AI agent to be aligned with traditional NLP relation extraction tasks with a structured output containing several sanity check mechanisms and post-processing guardrails. Structuring data in the JSON format allows us to quickly load and parse key pieces of data following a defined schema, including the drug and gene in the curated interaction. By employing normalization methods to these fields after AI agent curation, we are able to recognize and throw out extracted drugs or genes that do not harmonize to an existing biomedical concept, thus preventing incorrect or erroneous data. Further, by including an evidence field containing the string from the text that justified the extracted content, we can quickly sanity check whether an extracted entity existed in the literature or not. Lastly and notably, our extracted results were all performed with the commercially available Claude Opus 3.5 model. At the time this manuscript was completed, this model, while effective, was no longer top-of-the-line amongst even non-commercial LLM agents, leading us to believe using future iterations of the AI model may yield even better results.

Interestingly, our results highlight a high novelty rate of interactions from the literature. This high novelty could be viewed as a potential issue, as it may raise concerns about false positives and inaccuracies in interaction extraction. We instead argue that this is a desired feature and instead likely reflects limitations in the manual literature curation process (e.g. lag time from human curation to identify knowledge from newer literature).

Biocuration is resource-intensive, requiring substantial time investment from PhD-level biologists with expertise in the representation of computational knowledge^7^. Despite this, the benefits conferred from turning literature knowledge into structured knowledge are worth the investment, providing high-quality reviewed knowledge for high-throughput reuse by the scientific community as a whole. Thus, dgiLIT’s ability to identify novel interactions represents an important step towards the rapid transformation of published knowledge into structured literature.

However, our study does have limitations. Firstly, the initial search methods are highly dependent on the richness of available literature of any particular gene. For example, our results were highly positive for BCL2, a gene that is a hallmark of and frequently discussed for B-cell lymphomas and other cancers and is extensively studied in the literature both for experimental and basic research, and ongoing clinical trials. The effectiveness of dgiLIT for lesser described genes remains to be seen. While a tool that enables us to find new interactions for oncology and well described genes is valuable, the ability to extend that functionality to niche or lesser-publicized genes would further enhance value and biomedical knowledge. Secondly, while our tool speeds up the discovery time for biocuration, it does not remove the time barrier of human validation. The positive results with our BCL2 stress test could not have been assessed without manual work done by a human to review and assert that what was extracted was both real and valid. If data from this tool is to be included as a part of DGIdb, even as a new layer of searchable, AI-extracted data, validation by a human expert will still be required to ensure quality and trust of data.

Despite these limitations, our study highlights a new pipeline that has potential for greatly accelerating knowledge discovery efforts for new interactions in DGIdb. By performing a two step prioritization and AI curation task with guardrails, dgiLIT quickly and efficiently uncovers new interaction hypotheses for inclusion in our resource. Further, in cases where interactions are not novel, dgiLIT allows us to link new literature with old hypotheses, strengthening the existing data provenance, where available. Taken together, our results highlight a potential new mechanism for interaction knowledge discovery and a potential new layer of searchable, high quality data for our users.

## METHODOLOGY

### Code Availability

All code and example Jupyter notebooks for dgiLIT can be found in our public github repository located at https://github.com/dgidb/dgiLIT.

### Retrieval of Literature from PubTator3 annotations

The PubTator3 annotation set was retrieved from the FTP downloads section of the PubTator3 resource (available from https://ftp.ncbi.nlm.nih.gov/pub/lu/PubTator3/). The *gene2pubtator3* file specifically was used during the literature prioritization step to identify indexed literature containing references to designated genes of interest. All PMIDs containing a direct reference to a gene of interest were obtained and fed to the Entrez Programming Utilities (E-utilities) API to retrieve the complete set of published literature for downstream processing. While the PubTator3 data set does include automatically extracted relations as well, these annotations do not provide contextual evidence required for downstream curation and integration into DGIdb. Further, abstracts are needed for downstream processing via a generative language model in order to make use of context and output fields specific to DGIdb, such as interaction types and direct quoted evidence from the text.

### Lemmatization Groups

Lemma groups were constructed to capture “interaction words” in literature. Five broad categories of interaction words were created from interaction types encountered frequently in DGIdb: *Gene Regulation, Ligand Binding, Modulation, Sensitivity/Resistance, Pharmacogenomic Signals* (**Figure 2**). We curated interaction words associated with each parent category. These terms were grouped under the corresponding parent category and mapped to standardized ontology terms via the EMBL-EBI Ontology Lookup Service.. Lemma groupings and ontology definitions for interaction lemmas are provided in **Supplementary Table 1**.

### Subsetting of literature via bag-of-words for interaction lemmas

Abstracts of interest (as subset via PubTator3) were subject to a lemmatization process to identify the count of interaction words from our corresponding lemma groupings. Abstract text is lemmatized via the spaCy toolkit (e.g. “interacting” becomes “interact”) and exact matches with lemma groups are counted in a bag-of-words approach. The result of this count is used to perform a descending sort with the abstracts with the highest counts at the top of the list. This sort is thus used to prioritize literature containing the highest number of interaction words for downstream AI curation over those with less interaction words.

### AI Curation of Interactions from literature subsets

The top 100 abstracts that have been subset and prioritized (via PubTator3 and lemmatization, respectively) are further processed for downstream AI curation of drug-gene interactions. Named entity recognition (NER) is performed on text from the abstract to pre-identify drugs using a publically-available instance of BioBERT fine-tuned for NER of chemical entities^40^. PubTator3 chemical annotations were too noisy to be used directly thus identified chemical entities were normalized via the VICC therapy normalization tool^30^ to ensure only valid, ontologically grounded entities are included with the AI curation task. While BioBERT effectively captures chemical mentions from the text, the VICC therapy normalization tool contains curated groupings of aggregated identifiers and synonyms from public sources (including ChEMBL, DrugBank, RxNorm, IUPHAR/BPS, Drugs@FDA, ChemIDplus, Wikidata, etc.). Because of these differences in coverage, drugs identified via the NER output may not be present as an identifier in the harmonized groupings.

After pre-processing is complete, prompts are created programmatically for each abstract prior to the bulk AI curation task. The basic prompt template utilized contains a role description (e.g. scientific curator), a task description (e.g. curate drug-gene interactions from literature), a toolset description (e.g. definition of an interaction), and a sample output description (e.g. JSON template). A full length prompt is created for each abstract by filling in the basic prompt template with NER-identified therapeutics to consider (as part of the toolset) and the context on which to perform the task (the abstract text and PMID). Additionally, as part of the JSON output, a sanity check field is provided for the AI agent to confirm that the interaction is indeed occurring with a gene. Each full length prompt is saved as part of the output for data providence.

All assembled prompts are submitted to the AI client programmatically for curation (**Figure 3**). While dgiLIT can be used in a plug-and-play manner with any AI agent, for this report we utilized the commercially-available Claude Sonnet 3.5 model. For each abstract, full prompts were submitted via boto3. The model temperature parameter was set to 0.0 to reduce randomness of responses. Raw responses were obtained and parsed for JSON objects matching the supplied prompt template. Curated drug-gene interactions, interaction types, and evidence statements were extracted from JSON objects and subjected to downstream processing for data quality. Curated drug and gene entities were normalized via the VICC therapy and gene normalization services, respectively. Entities that did not normalize to a grounded drug or gene were thrown out. Similarly, curated evidence that failed the sanity check statement was thrown out.

### Quantification of Publication Statistics

Publication statistics and associated metadata for PMIDs obtained as part of the curated abstract set were obtained, including the SCImago Journal Rank (SJR) indicator, the journal h-index, and associated Medical Subject Heading (MeSH) terms. SJR indicators and journal h-indices were obtained from the SCImago Journal & Country Rank public dataset (available from <u>scimagojr.com</u>). MeSH headings for each PMID were programmatically obtained using the Entrez Programming Utilities (e-utilities) API suite.

### Quantification of Ontological Expansion

Ontological expansion was assessed by comparing the set of normalized entities from the curated interactions against the entities already present inside DGIdb. Entities that were not present in DGIdb during the fall 2024 update but were present in the curated interaction set were tallied as expansion, even though this reflects knowledge that was added to DGIdb in the interim between our study start and conclusion.

### Quantification of Novel Interactions for DGIdb

Drug-gene interaction pairs were assembled from the AI Curation results. Genes from each pair were queried against the existing DGIdb database with the resulting search being interrogated for the matching drug. When an interaction is found in the DGIdb database, the number of sources supplying an interaction claim is recorded. Queries with zero results correspond to a novel interaction.

### Human Expert Evaluation of AI curated Interactions

The full set of AI-curated interactions, evidence statements, interaction directionality, and PMIDs were supplied to an expert human biomedical curator. The curator was asked to evaluate the AI-curated interactions, specifically to determine whether the drug, gene, and evidence statements were real and appeared in the text. Next, the curator was asked to validate the AI supplied directionality of the interaction (e.g. inhibiting or activating) and whether that could be assessed from the abstract alone or if the full-text was needed for additional support. Directionality was determined by matching the curated interaction to a supported interaction type(available from https://dgidb.org/about/overview/types-and-directionality). Additionally, the curator evaluated the AI curated evidence statements by either verifying them as the most representative statements from the text for the interaction, or if not sufficient, selecting a more appropriate quote directly from the text.

Lastly, the curator assessed the confidence in interaction directionality, with lower scores indicating lower confidence. Scores were defined as follows: 1) low confidence: inferred interactions lacking clear evidence of an interaction-type match; 2) moderate confidence: interactions matching multiple potential interaction types; and 3) high confidence: interactions with a clear, well-supported interaction type. The responses for validity assessment, depth required for validation, and interaction directionality were collected and quantified. Assessment data is available for review in **Supplemental Table 4**.

**Supplementary Figure 1.**
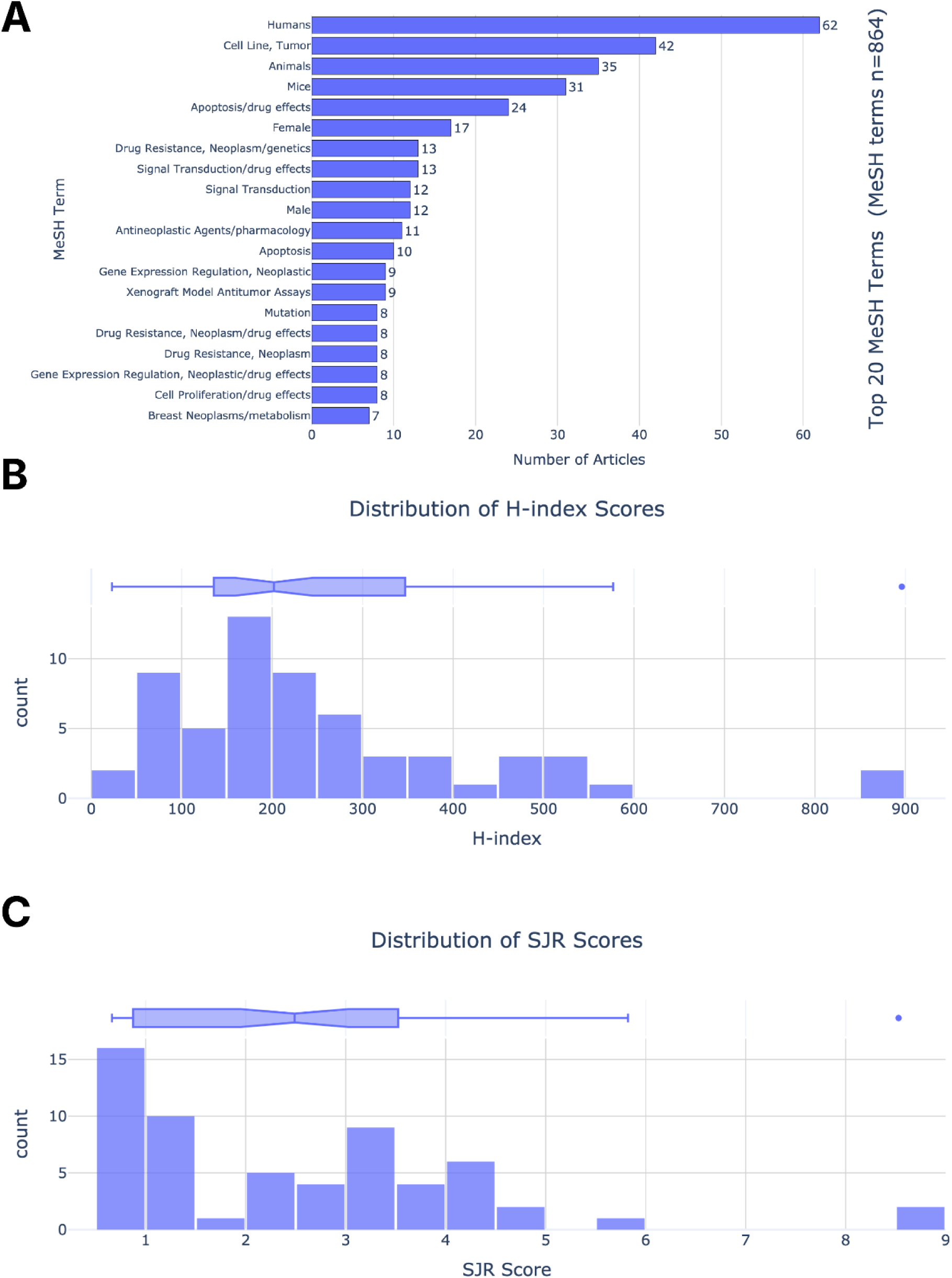
Classification, Citation, and Impact Metrics for PMIDs used for AI Curation Task. **(A)** Articles selected via dgiLIT demonstrated strong relevance to translational medicine as indicated via MeSH terms. **(B-C)** Articles selected via dgiLIT showed a high impact with a high average journal article H-index and SJR score.

**Supplementary Figure 2.**
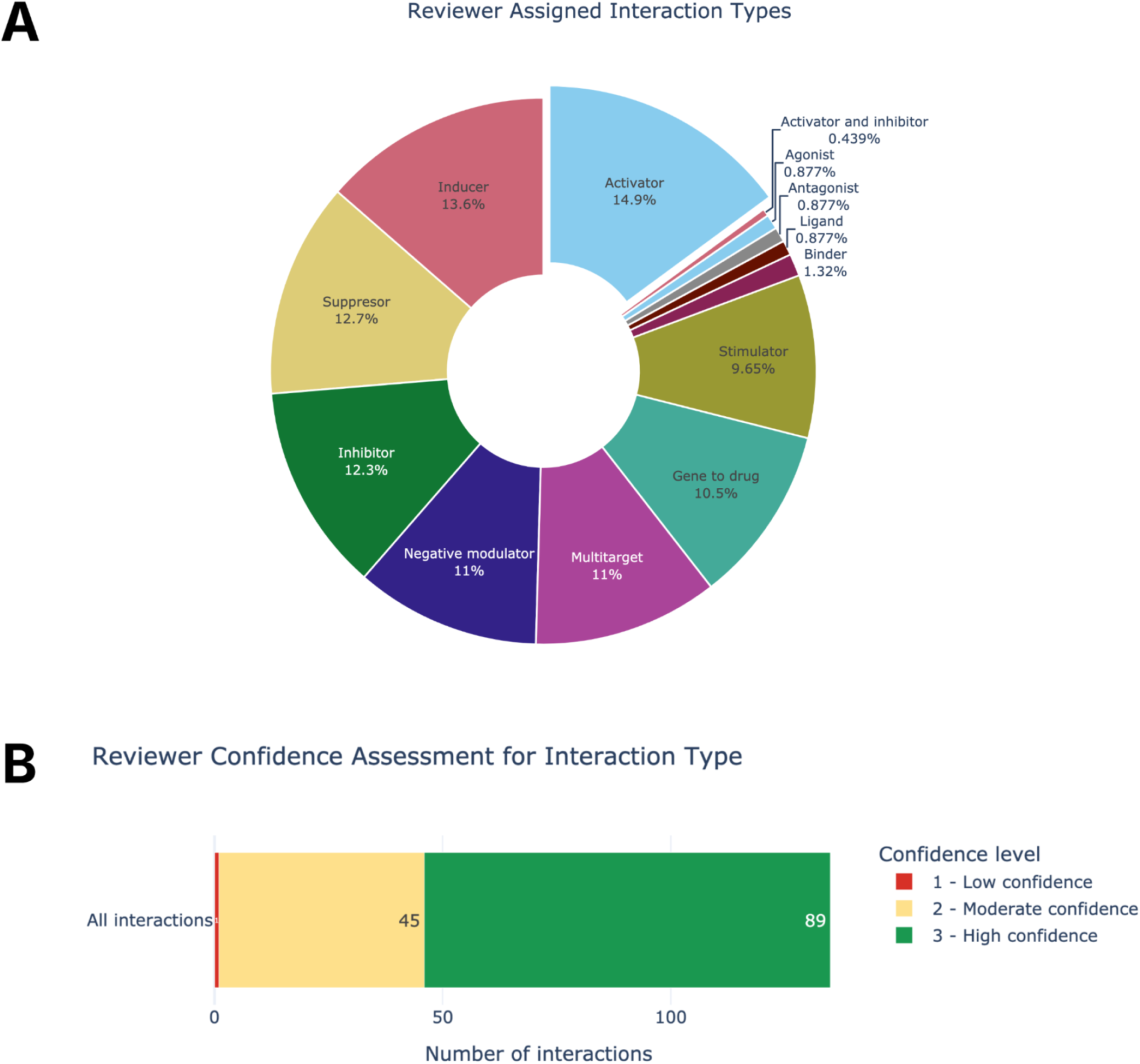
**Reviewer Assessment of Interaction Types from AI Curated Interactions**. (**A**) The human expert reviewer was asked to assess the likely interaction type for AI curated interactions (using type definitions available from https://dgidb.org/about/overview/types-and-directionality). (**B**) The human expert reviewer was asked to provide a confidence value for their interaction type assignment using the following scale: 1) low confidence: inferred interactions lacking clear evidence of an interaction-type match; 2) moderate confidence: interactions matching multiple potential interaction types; and 3) high confidence: interactions with a clear, well-supported interaction type.

## Notes

### Competing Interest Statement

The authors have declared no competing interest.

### Funding

- National Human Genome Research Institute: R00HG010157

